# Hybrid systems modeling of ecological population dynamics

**DOI:** 10.1101/2020.03.28.013524

**Authors:** Abhyudai Singh, Brooks Emerick

## Abstract

Discrete-time models are the traditional approach for capturing population dynamics of insects living in the temperate regions of the world. These models are characterized by an update function that connects the population densities from one year to the next. We revisit classical discrete-time models used for modeling interactions between two insect species (a host and a parasitoid), and provide novel result on the stability of the population dynamics. In particular, for a class of models we show that the fixed point is stable, if and only if, the host equilibrium density is an increasing function of the host’s reproduction rate. We also introduce a hybrid approach for obtaining the update functions by solving ordinary differential equations that mechanistically capture the ecological interactions between the host and the parasitoid. This hybrid approach is used to study the suppression of host density by a parasitoid. Our analysis shows that when the parasitoid attacks the host at a constant rate, then the host density cannot by suppressed beyond a certain point without making the population dynamics unstable. In contrast, when the parasitoid’s attack rate increases with increasing host density, then the host population density can be suppressed to arbitrarily low levels. These results have important implications for biological control where a natural enemy, such as a parasitoid wasp, is introduced to eliminate a pest that is the host species for the parasitoid.

## I. Introduction

Insect population dynamics has been extensively studied using two different approaches: continuous-time and discrete-time models. The continuous-time framework is generally used to model populations with overlapping generations and all year-round reproduction. In contrast, discrete-time models are more suited for populations in temperate regions of the world that have non-overlapping generations and reproduce in a discrete pulse determined by season [1].

Here we revisit discrete-time models capturing the antagonistic interaction between two insect species (a host and a parasitoid). A typical life cycle of the host and the parasitoid is shown in Fig. 1, and consists of adult and hosts emerging during spring, laying eggs that hatch into larvae. Host larvae then overwinter in the pupal stage, and metamorphosize as adults the following year. As adult hosts die after laying eggs, there is no overlap between generations across years. The host becomes vulnerable to attacks by a parasitoid wasp at one stage of its life cycle. For the sake of convenience, we assume the host’s vulnerable stage to be the larval stage. Adult female parasitoids emerge during spring, search and attack hosts by laying an egg into the body of the host. While adult parasitoids die after this time window, the parasitoid egg hatches into a juvenile parasitoid that grows at the host’s expense by using it as a food source, and this ultimately results in the death of the host. The juvenile parasitoids pupate, overwinter, and emerge as adult parasitoids the following year. There are more than 65,000 different species of parasitoid wasps and we refer the reader to [2]–[4] for fascinating details on parasitoid classification, life history and behavior^1^. Synchronized life cycles, with no overlap of generations in both the host and the parasitoid makes discrete-time models highly appropriate for these systems.

**Fig. 1:**
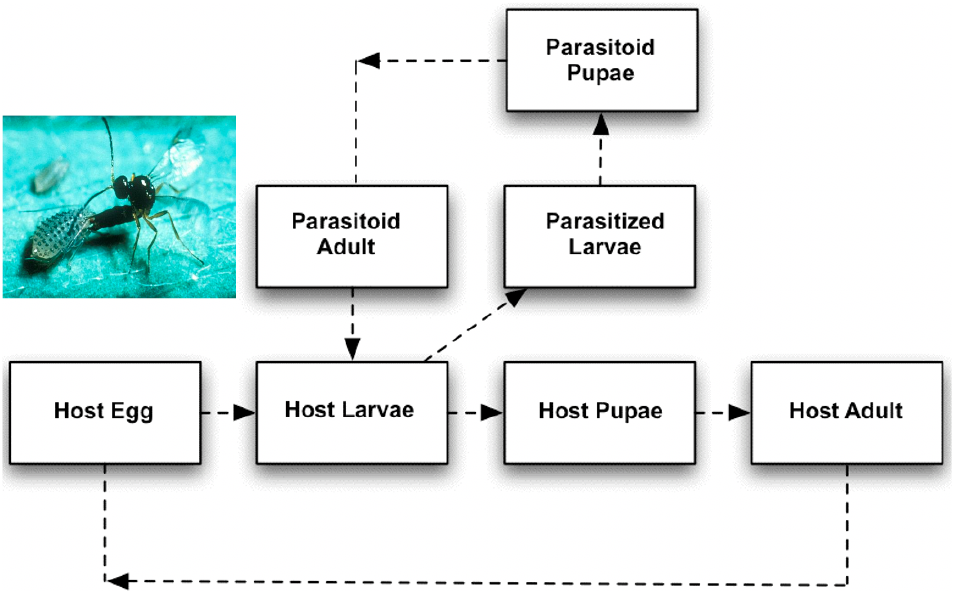
Life cycle of the host and the parasitoid. Inset shows the picture of a parasitoid wasp laying an egg into the body of its host (spotted alfalfa aphid). Picture taken from https://en.wikipedia.org/wiki/Parasitoid.

We provide novel results on the stability of a general class of discrete-time models describing host-parasitoid interactions. Following a rich tradition of using hybrid or semi-discrete approaches to model ecological population dynamics [5]–[10], we discuss a hybrid formulation that solves a nonlinear different equation to obtain discretetime models for host-parasitoid systems. Importantly, this mechanistic derivation of models using the hybrid approach can provide qualitatively different results compared to earlier phenomenological approaches. Finally, in the context of biological control of pests, we investigate different forms of parasitoid attack rates that lead to efficient suppression of the host population.

## II. Model Formulation

A model describing host-parasitoid d discrete-time is given by

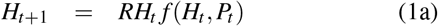

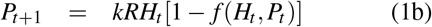

where *H*_*t*_ and *P*_*t*_ are the adult host, and the adult parasitoid densities, respectively, at the start of year *t* [11]–[13]. If the host is vulnerable to the parasitoid at its larval stage, then *RH*_*t*_ is the host larval density exposed to parasitoid attacks at the start of the vulnerable stage, where *R*>1 denotes the number of viable eggs produced by each adult host. Parasitoids attack host larvae during the vulnerable period leading to two populations of hosts: parasitized and un-parasitized larvae. The function *f*(*H*_*t*_, *P*_*t*_) < 1 is the fraction of host larvae escaping parasitism and can depend on both *H*_*t*_ and *P*_*t*_. We assume *f* to be a continuously differentiable function in both arguments and refer to it as the *escape response*. In the absence of the parasitoid

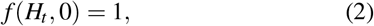

and the host population grows unboundedly as

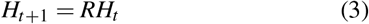

where *R* can be interpreted as the host’s growth rate. In (1), *RH*_*t*_*f*(*H*_*t*_, *P*_*t*_) is the total larval density escaping parasitism to become adult hosts for next year. Finally, *RH*_*t*_ [1 − *f*(*H*_*t*_, *P*_*t*_)] is the net density of parasitized larvae, with each larva giving rise to *k* adult female parasitoids in the next generation.

The simplest formulation of (1) is the classical Nicholson-Bailey model

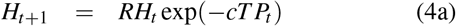

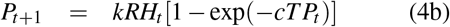

where *c* > 0 represents the rate at which parasitoids attack hosts, and *T* is the duration of the host vulnerable stage [14]. The model assumes that parasitoids search for hosts randomly, are never egg-limited, and have quick handling times. If the number of parasitoid attacks per host follows a Poisson distribution with mean *cTP*_*t*_, then the escape response *f*(*H*_*t*_, *P*_*t*_) = exp(−*cTP*_*t*_) is the probability of zero attacks in the Poisson distribution. In Section IV, we provide another mechanistic interpretation of the Nicholson-Bailey model. A typical time series of the Nicholson-Bailey model is shown in Fig. 2. Both populations grow at low densities, but at large host densities the parasitoid begins to overexploit the host. This leads to a crash in the host population, followed by a crash of the parasitoids. These cycles of overexploitation and crashes result in an unstable interaction with both populations exhibiting diverging oscillations. Interestingly, these diverging oscillations have been recapitulated in several laboratory studies of specific host-parasitoid species providing some model validation, at least in controlled lab settings.

**Fig. 2:**
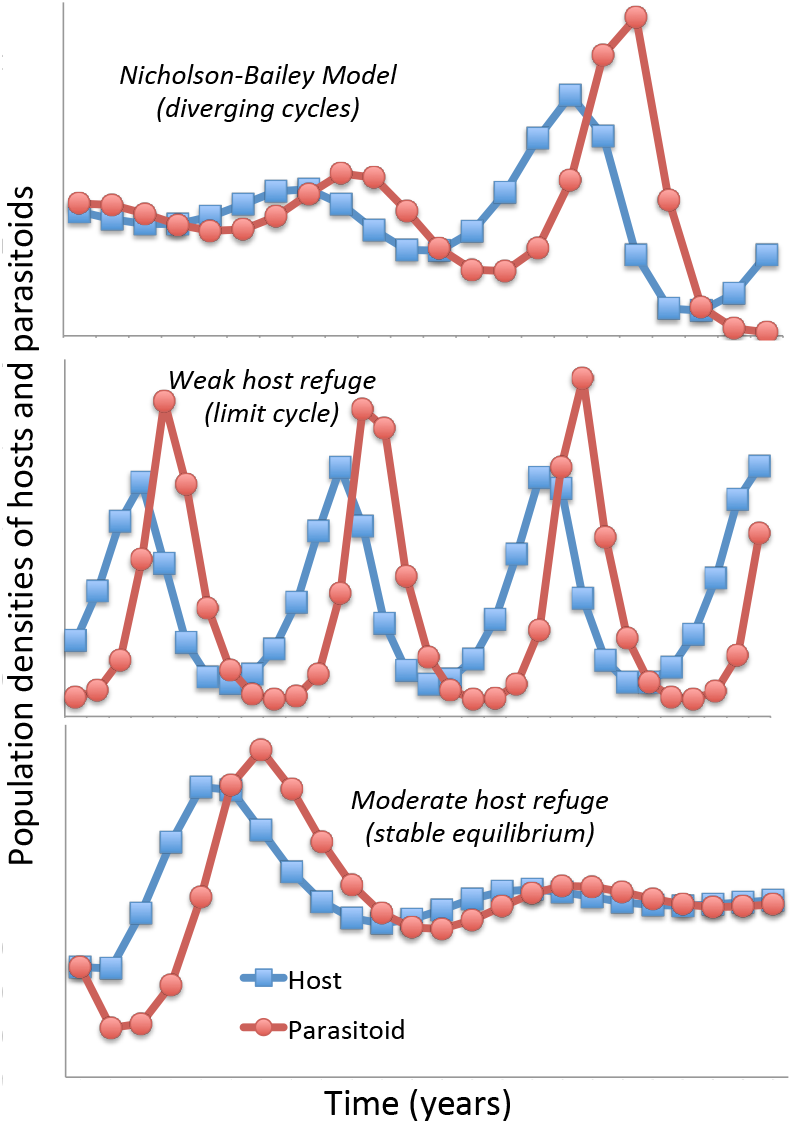
A typical host-parasitoid population time series for the Nicholson-Bailey model (top), model with weak host refuge (*μ* = 0.05 in (5); middle) and moderate host refuge (*μ* = 0.40 in (5); bottom). Host reproduction rate assumed to be *R* = 2, *T* = 1 and *k* = 1.

Several mechanisms have been proposed in literature that can stabilize the host-parasitoid population dynamics. One such mechanism is a host refuge where a constant fraction of hosts are protected from parasitism. A constant host fraction *μ* in the refuge leads to the following model

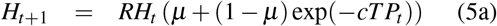

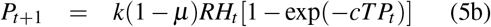

[1]. While a weak-refuge results in bounded oscillations, a moderate-refuge stabilizes the population dynamics (Fig. 2). However, stability is again lost for a strong-refuge with both hosts and parasitoids growing unboundedly. Next, we provide some novel results on the stability of the discrete-time model (1).

## III. General Stability Analysis

The fixed point of the discrete-time model (1) are the population densities that remain constant across years. Substituting *P*_*t*+1_ = *P*_*t*_ = *P\** and *H*_*t*+1_ = *H*_*t*_ = *H\** in (1) shows that apart from the trial fixed point *P\** = *H\** = 0, the non-trivial equilibrium is the solution to the following equations

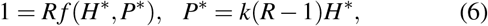

where *H\** and *P\** denote the host and parasitoid densities at equilibrium, respectively.

Solving these equations for the Nicholson-Bailey model yields a single non-trivial equilibrium point

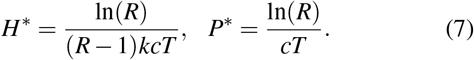

Note that the host equilibrium levels decrease, while the parasitoid equilibrium levels increase, with increasing host growth rate *R*.

Considering small perturbations *h*_*t*_ and *p*_*t*_

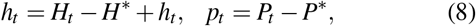

and linearizing model nonlinearities in (1) around the equilibrium, results in the following *linear* discrete-time system

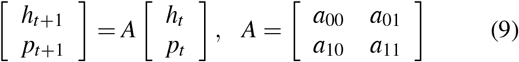

where the entries of the Jacobian matrix *A* are given by

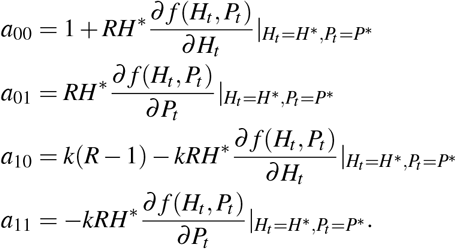

Here 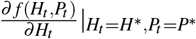 represents the partial derivative of the escape response with respect to the host density evaluated at the equilibrium point.

Standard stability analysis shows that the non-trivial equilibrium point is stable, if and only if, the following three conditions hold

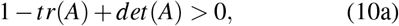

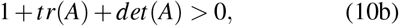

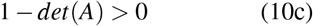

where the trace and determinant of the *A* matrix are

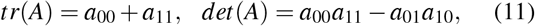

respectively [15]. The Theorem below shows that in some cases these stability conditions can be remarkably simplified.

### Theorem

Consider the discrete-time model

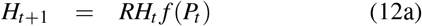

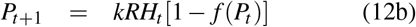

where the escape response *f*(*P*_*t*_) is a monotonically decreasing function of the parasitoid density only, with *f*(0) = 1. Then, the model has a unique non-trivial fixed point given by the solution to the equation

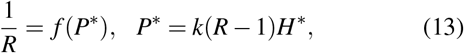

and this fixed point is stable, if and only if,

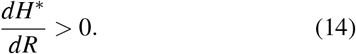

We refer the reader to the appendix for a proof of the theorem. Briefly, if the escape response *f*(*P*_*t*_) only depends on parasitoid density then the first two conditions in (10) always hold, and stability is completely determined by the third inequality 1 − *det*(*A*) > 0 which can be rewritten as (14). Recall that *H\** in the Nicholson-Bailey model is decreasing with *R*, and its instability is reflective of this simplified stability criterion. Applying the theorem to the host refuge model (5) we obtain the following equilibrium host density

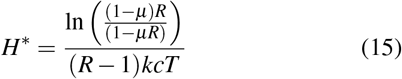

that is only defined for *μ* < 1/*R*. Fig. 3 plots *H*\* as a function of *R* for different values of the refuge fraction *μ*. The plots shows that initially *H\** decreases with increasing *R* (hence, an unstable equilibrium), but beyond a critical host growth rate, *H\** increases with increasing *R* leading to stability (as seen in Fig. 2). For example, when *μ* = 0.4, the population dynamics is stabilized for *R* > 1.375.

**Fig. 3:**
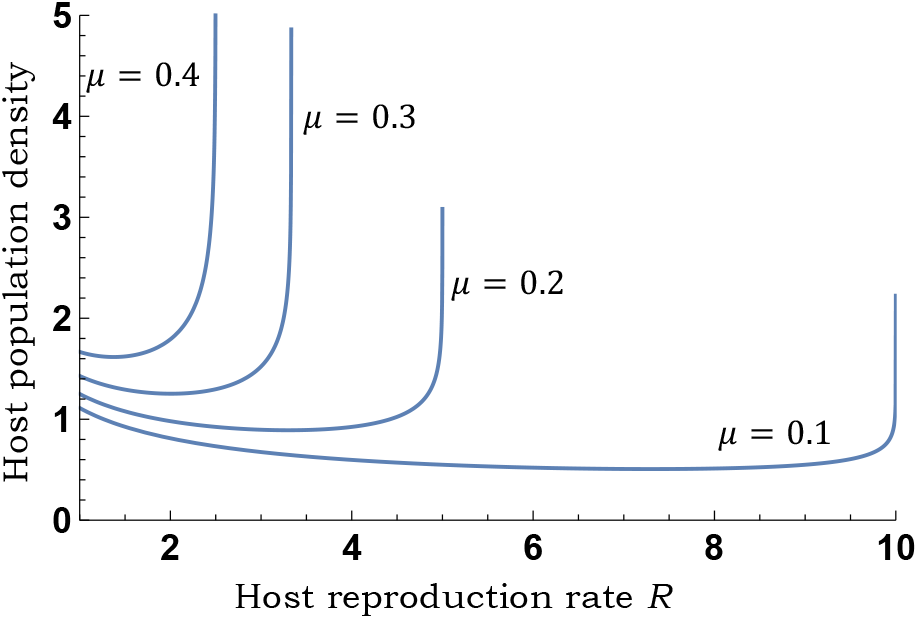
Host equilibrium density (15) for the refuge model (5) as a function of the growth rate *R*. For a given *μ*, *H\** first decreases with increasing *R* (unstable equilibrium), but beyond a critical host growth rate, *H\** increases with increasing *R* (stable equilibrium).

Another mechanism know to stabilize host-parasitoid interactions is when the parasitoid’s attack rate *c* in (4) decreases with increasing parasitoid density due to interference between parasitoids. Rewriting 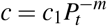, where *c*_1_ > 0 and 0 < *m* < 1 results in the following discrete-time model

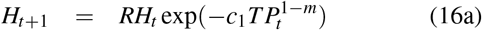

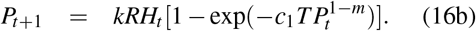

A straightforward application of the theorem shows that the model (16) is stable for sufficiently large value of *m* given by

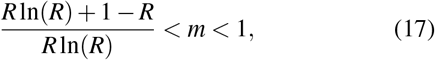

which corresponds to 0.28 < *m* < 1 for *R* = 2, and 0.5 < *m* < 1 for *R* = 5.

Finally, using the fact that *P\** = *k*(*R* − 1)*H\**, the above condition on *H\** can also be written in terms of the parasitoid equilibrium density

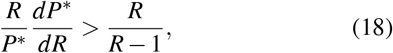

revealing that stability requires parasitoid densities to increase sufficiently fast with increasing *R*.

It is important to emphasize that condition (14) can only to be used for a host-independent escape response. The above theorem begs the question: can this simple stability criterion be generalized when *f*(*H*_*t*_, *P*_*t*_) depends on both populations? It turns out that in this case, the stability conditions (10) can be graphically represented in terms of two biologically-relevant quantities:

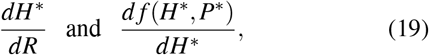

where the latter denotes the sensitivity of the escape response to the host density. Fig. 4 shows that stability arises in a narrow band, and is more likely to occur when the escape response is a decreasing function of the host density, rather than an increasing function. An interesting insight from this figure is that stability can arise in the Nicholson-Bailey model from two orthogonal mechanisms: a host refuge that moves the point upwards into the stability region; alternatively a Type-III functional response that moves the point to the left into the stability region. We discuss Type III functional response in more detail in the next section.

**Fig. 4:**
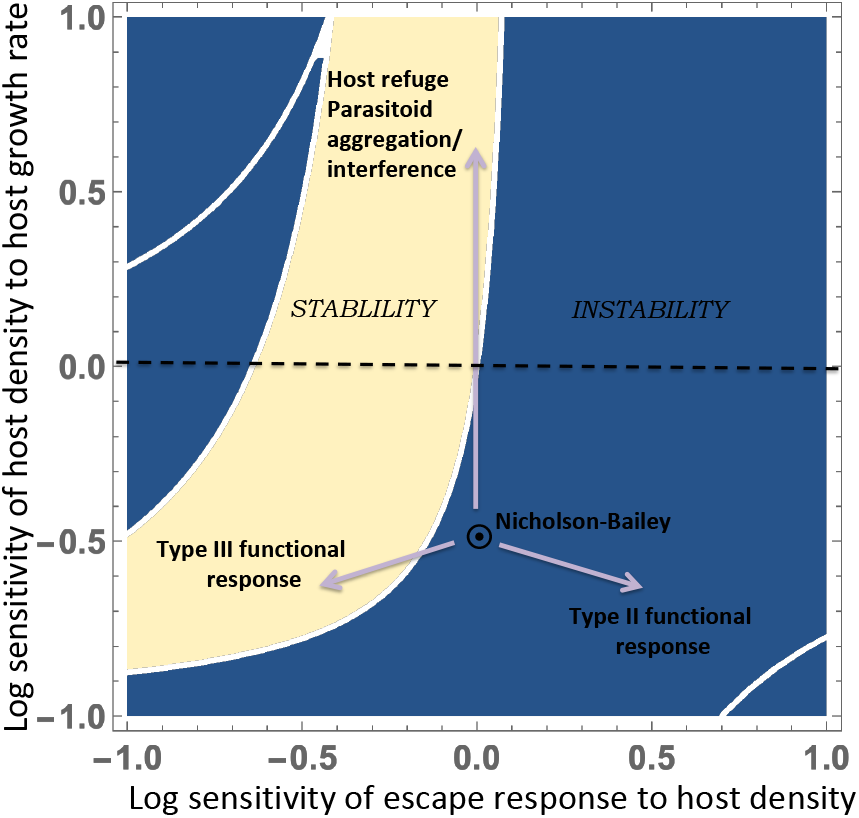
Stability regions for the host-parasitoid model (1) in terms of the sensitivity of the host density to the host growth rate *dH*/dR*, and the sensitivity of the escape response to the host density *d f /dH\**. The Nicholson-Bailey model corresponds to *dH*/dR* < 0 and *d f /dH\** = 0 and is unstable. Stability arises in two orthogonal ways: 1) an increase in *dH*/dR* to make it positive, which happens with parasitoid interference/aggregation or host refuge; 2) a decrease in *d f /dH\** which happens with a Type III functional response in the parasitoid attack rate. For this figure, the host growth rate is assumed to *R* = 2, *k* = 1 and the stability region shrinks with increasing *R*.

## IV. Hybrid formulation of update functions

For a majority of host-parasitoid models the escape response *f* in (1) is phenomenologically chosen or designed to recapitulate field observations. Recent work has proposed a mechanistic hybrid framework for deriving the discrete-time model, where ordinary differential equations are used to track population densities within the host vulnerable period of a given year [16]–[18]. The solution of the differential equations at the end of the vulnerable period predicts the population densities for next year. We discuss this semi-discrete formulation in further detail below.

Let *τ* denote the time within the host vulnerable stage that varies from 0 to *T* corresponding to the start and end of the vulnerable stage. The densities of parasitoids, unparasitized and parasitized host larvae at time *τ* within the vulnerable stage of year *t* are represented by *P*(*τ, t*), *L*(*τ, t*), *I*(*τ, t*), respectively. These densities evolve as per the dynamical system

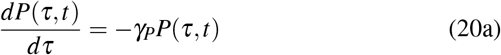

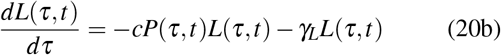

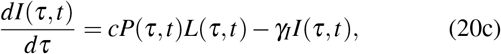

where *c* represent the parasitoid’s attack rate *per host*, and *γ*_*P*_, *γ*_*L*_, *γ*_*I*_ are the death rates of the respective species. Assuming *P*_*t*_ parasitoids, *RH*_*t*_ host larvae, and no parasitized larvae at the start of the vulnerable period (*τ* = 0), solving the above differential equations with initial conditions

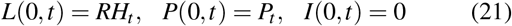

predicts the parasitized and unparasitized larval population at the end of the season (*τ* = *T*). This leads to a more general discrete-time model

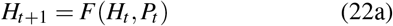

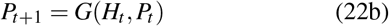

where update functions are obtained by setting

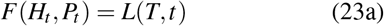

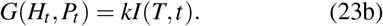

Solving (20) for a constant attack rate *c* with no mortalities (*γ*_*P*_ = *γ*_*L*_ = *γ*_*I*_ = 0) yields

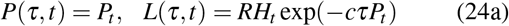

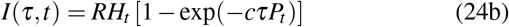

which using (23) yields the Nicholson-Bailey model (4).

Interestingly, it turns out that discrete-time models formulated using the hybrid approach can give qualitatively different results than phenomenologically designed updated functions. This point was illustrated in [16] for the scenario when the attack rate *c* becomes dependent on the density of un-parasitized hosts

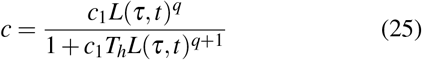

for positive constants *c*_1_, *q* and *T*_*h*_, In ecological literature, the case of *q* = 0 has been referred to as a Type II functional response where

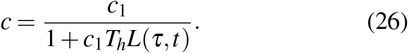

When *q* = 0, the parasitoid attack rate across all hosts takes the form

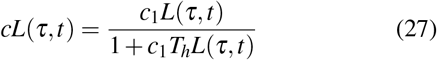

which saturate to

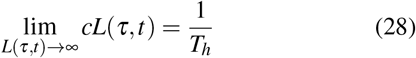

at high host densities, where the handling time *T*_*h*_ can be interpreted as the average time taken by a parasitoid to finish parasitizing a host. The case of *q* ≥ 1 is referred to as a Type III functional response where *cL*(*τ, t*) becomes a sigmoidal function of *L*(*τ, t*). Setting a host-dependent attack rate (25) in (20) results in a stable discrete-time model for *q* > 1 assuming that the handling time is significantly shorter than the vulnerable stage duration (*T*_*h*_ ≪ *T*) [16]. Interestingly, the host equilibrium here is still a decreasing function of the host growth rate (as in the Nicholson-Bailey model), but a Type III response makes the fraction of hosts escaping parasitism a decreasing function of the host density. In the context of Fig. 4, this corresponds to shifting the Nicholson-Bailey point to the left into the stability region. In contrast, a Type II response (*q* = 0) moves the point to the right to further destabilize the population dynamics.

Several prior works incorporated functional responses phenomenologically by simply substituting

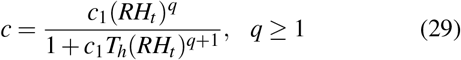

in the Nicholson-Bailey model (4), and the resulting discrete-time model is unstable for all values of *q* [19]. In summary, a Type III functional response (*q* > 1) stabilizes the population interaction in the mechanistically formulated hybrid approach, but not in the phenomenological approach.

## V. Limits of host suppression by parasitoids

We next use the hybrid formulation to investigate the suppression of host population density by a parasitoid. As done in earlier models [20]–[22], we first consider a self-limitation in the hosts ability to grow (due to finite food resources or attacks by other nature enemies) by considering a host mortality rate *γ*_*L*_ = *c*_*h*_*L*(*τ, t*) in (20) that is proportional to the host density. For a constant parasitoid attack rate *c* and *γ*_*P*_ = *γ*_*I*_ = 0, the above hybrid approach yields

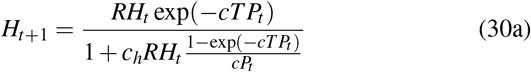

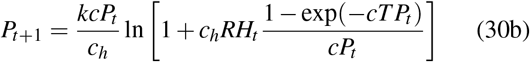

[16]. As expected, in the limit *c*_*h*_ → 0 this model converges back to the Nicholson-Bailey model. In the absence of parasitoids, this leads to

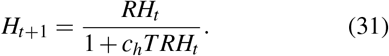

which has a stable fixed point

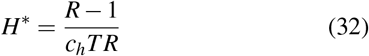

representing the host population before the introduction of the parasitoid. In addition to this no-parasitoid equilibrium, the other fixed point of (30) is given by

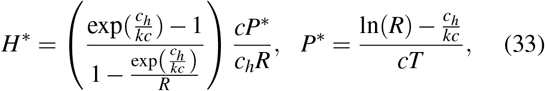

and is stable for

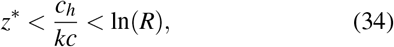

where *z\** is the solution to

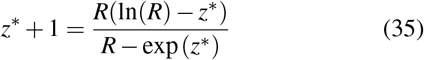

[16]. The quantity

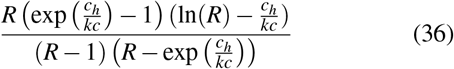

represents the ratio of the host equilibrium with parasitoids and without parasitoids, as given by (33) and (32), respectively. As one varies *c*_*h*_/*c* in the stability region (34), the maximum host suppression while ensuring the stability of the fixed point (33) is given by

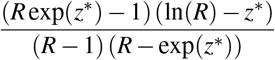

and is computed to be 0.34 and 0.29 for *R* = 2 and 10, respectively. Thus, for a constant parasitoid attack rate, the host population density cannot be suppressed more than ≈30% below its density in the absence of the parasitoid. Can host suppression be more efficient with a density-dependent parasitoid attack rate?

To address this question, we consider a Type III functional response where the attack rate takes the form (25). For simplicity we consider the case of *q* = 1 and fast handling times (*T*_*h*_ → 0) which reduce (25) to

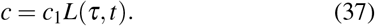

In essence, the ability of a parasitoid to locate, attack and parasitize a host, increases with increasing host density. Using (37), *γ*_*L*_ = *c*_*h*_*L*(*τ, t*), *γ*_*P*_ = *γ*_*I*_ = 0 in (20) yields

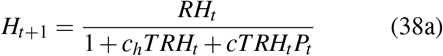

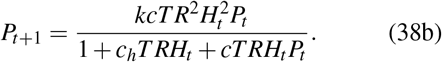

The model again has two non-trivial equilibriums. The first is when the parasitoid is absent, and the host density converges to (32). The second fixed point where both species coexist is given by

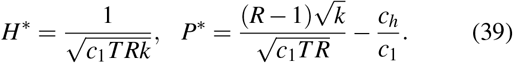

Stability analysis as outlined in Section III shows that (39) is stable as long as

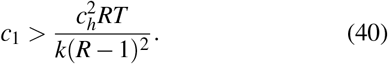

This implies that the ratio of host density with and without parasitoids

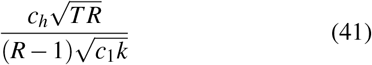

can be made arbitrarily small with increasing attack rate parameter *c*_1_ but with a dependence that scales as 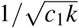. This point is illustrated in Fig. 5 where the time evolution of host densities are shown for increasing values of *c*_1_.

**Fig. 5:**
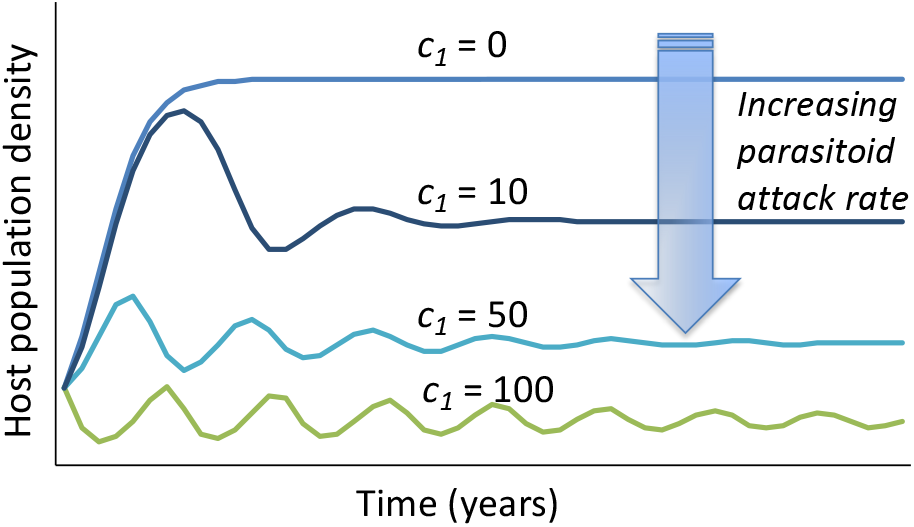
Plots of host population density *H*_*t*_ as a function of time *t* for model (38). With increasing parasitoid attack rate parameter *c*_1_ in (37), the host density can be suppressed to arbitrary low levels while ensuring population stability. Other parameters taken as *R* = 2, *c*_*h*_ = 1, *T* = 1 and *k* = 1.

## VI. Conclusion

In this contribution, we have focused on the ecological interaction between hosts and parasitoids that holds tremendous potential in biological control of pests [23]–[28]. We provide novel result on the stability of host-parasitoid interactions modeled as per (1), and characterized by an escape response *f*(*H*_*t*_, *P*_*t*_). The result is graphically illustrated in Fig. 4 in terms of two quantities: the rate at which the host equilibrium changes with the hosts growth rate (*dH*/dR*), and the sensitivity of the escape response to the host density at equilibrium (*d f /dH*). For several models, such as (4), (5), (16) the escape response is invariant of the host density, in which case the stability reduces to the host density being an increasing function of the host growth rate. Another key insight that can be tested with field observations is that stability is more likely to emerge when the escape response is a decreasing function of the host density (i.e., a lager fraction of hosts do not get parasitized at higher host densities), than an increasing function.

We also introduced a hybrid approach for deriving the discrete-time model by solving a set of nonlinear differential equations describing the population dynamics during the host’s vulnerable stage. The hybrid approach was used to study the limit of host suppression while still keeping the host-parasitoid interaction stable. Our prior work had shown that for a constant parasitoid attack rate, the host cannot be suppressed beyond a certain limit [16]. Interestingly, our analysis here shows that an attack rate proportional to the host density can lead to arbitrary low host suppression (Fig. 5). In future work, we will investigate host suppression with other stabilizing mechanisms, such as, host refuge, variation of risk across the host population [13], [29]–[31], and parasitoid aggregation [32]–[34]

Another aspect to explore using the hybrid formulation is spatial dynamics where host/parasitoid can move between patches [35]–[38]. Our preliminary work on this topic has been quite promising and shows that density-dependent movement between patches can have a stabilizing effect on the population dynamics [39].

## Appendix

Consider the discrete-time model (12) whose unique non-trivial fixed point is given by

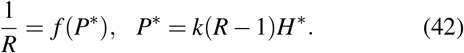

Denoting small fluctuation around the equilibrium *H\** and *P\** by *h*_*t*_ := *H*_*t*_ − *H\** and *p*_*t*_ := *P*_*t*_ − *P\**, respectively, one obtains using linearization the following linear discrete system

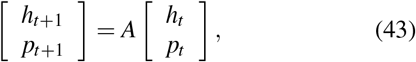

with

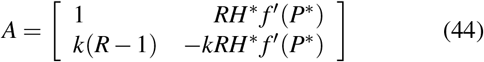

where

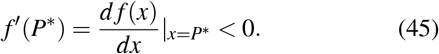

The stability conditions (10) applied to (44) shows the model fixed point is stable, if and only if, the following inequalities hold

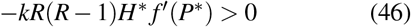

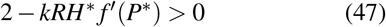

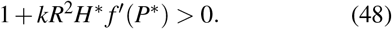

Given that *R* > 1 and the fact that *f′*(*P\**) < 0 (*f* is a monotonically decreasing function), inequalities (46)-(47) always hold and the stability condition is given by inequality (48), which using *P\** = *k*(*R−* − 1)*H\** becomes

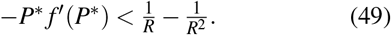

Differentiating the first equation in (42) with respect to *R* we have

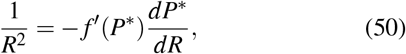

which using *P\** = *k*(*R−* 1)*H\** can be written as

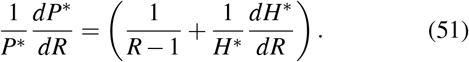

Substituting (51) in (50) gives us

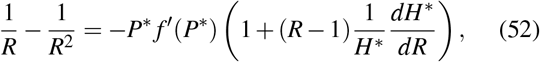

which using the stability condition (49) reduce to

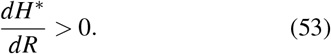

## Footnotes

1 Host-parasitoid interactions motivated the 1979 Hollywood movie *Alien* where a human is infected by an alien life-form. The alien develops within the human and later emerges killing the human.

## References

[1] W. W. Murdoch, C. J. Briggs, and R. M. Nisbet, Consumer-Resouse Dynamics. Princeton, NJ: Princeton University Press, 2003.

[2] A. E. Hajek, Insect parasitoids: attack by aliens. Cambridge University Press, 2004, p. 145169.

[3] H. C. J. Godfray, Parasitoids; Behavioral and Evolutionary Ecology. 41 William St, Princeton, NJ 08540: Princeton University Press, 1994.

[4] J. Waage and D. Greathead, Insect Parasitoids. Academic Press, 1986.

[5] E. Pachepsky, R. M. Nisbet, and W. W. Murdoch, “Between discrete and continuous: Consumer-resource dynamics with synchronized reproduction,” Submitted for publication, 2007.

[6] T. M. Eskola and S. A. Geritz, “On the derivation of discrete-time population models by varying within-season patterns of reproduction and aggression,” Bulletin of Mathematical Biology, vol. 69, pp. 329–346, 2007.

[7] C. J. Dugaw, A. Hastings, E. L. Preisser, and D. R. Strong, “Parasitoid-mediated effects: apparent competition and the persistence of host-parasitoid assemblages,” Bulletin of Mathematical Biology, vol. 66, no. 3, pp. 583–594, 2004.

[8] M. B. Bonsall and M. P. Hassell, “Parasitoid-mediated effects: apparent competition and the persistence of host-parasitoid assemblages,” Researches on Population Ecology, vol. 41, no. 1, pp. 59–68, 1999.

[9] C. J. Briggs and H. C. J. Godfray, “The dynamics of insect-pathogen interactions in seasonal environments,” Theoretical Population Biology, vol. 50, pp. 149–177, 1996.

[10] S. A. Geritz and E. Kisdi, “On the mechanistic underpinning of discrete-time population models with complex dynamics,” J. of Theoretical Biology, vol. 228, no. 2, pp. 261–269, 2004.

[11] M. P. Hassell. New York: Oxford University Press, 2000.

[12] W. S. C. Gurney and R. M. Nisbet, Ecological Dynamics. Oxford University Press, 1998.

[13] A. Singh, W. W. Murdoch, and R. M. Nisbet, “Skewed attacks, stability, and host suppression,” Ecology, vol. 90, no. 6, pp. 1679–1686, 2009.

[14] A. Nicholson and V. A. Bailey, “The balance of animal populations. part 1.” Proc. of Zoological Society of London, vol. 3, pp. 551–598, 1935.

[15] S. Elaydi, An Introduction to Difference Equations. Newyork: Springer, 1996.

[16] A. Singh and R. M. Nisbet, “Variation in risk in single-species discrete-time models,” Mathematical Biosciences and Engineering, vol. 5, pp. 859–875, 2008.

[17] B. K. Emerick and A. Singh, “The effects of host-feeding on stability of discrete-time host-parasitoid population dynamic models.” Mathematical Biosciences, vol. 272, pp. 54–63, 2016.

[18] A. Singh and R. M. Nisbet, “Semi-discrete host-parasitoid models,” J Theor Biol, vol. 247, no. 4, pp. 733–742, 2007.

[19] M. P. Hassell and H. N. Comins, “Sigmoid functional responses and population stability,” Theoretical Population Biology, vol. 14, pp. 62–66, 1978.

[20] M. G. Neubert and M. Kot, “The subcritical collapse of predator populations in discrete-time predator-prey models.” Mathematical Bioscience, vol. 110, pp. 45–66, 1992.

[21] M. P. Hassell and G. C. Varley, “New inductive population model for insect parasites and its bearing on biological control,” Nature, vol. 223, pp. 1133–1137, 1969.

[22] R. M. May, M. P. Hassell, R. M. Anderson, and D. W. Tonkyn, “Density dependence in host-parasitoid models,” J. of Animal Ecology, vol. 50, pp. 855–865, 1981.

[23] P. K. Abram, J. Brodeur, V. Burte, and G. Boivin, “Parasitoid-induced host egg abortion; an underappreciated component of biological control services provided by egg parasitoids.” Biological Control, no. 98, pp. 52–60, 2016.

[24] M. A. Jervis, B. A. Hawkin, and N. A. C. Kidd, “The usefulness of destructive host-feeding parasitoids in classical biological control: theory and observation conflict,” Ecological Entomology, vol. 21, no. 1, pp. 41–46, 1996.

[25] T. Ueno, “Selective host-feeding on parasitized hosts by the parasitoid itoplectis naranyae (hymenoptera: Ichneumonidae) and its implication for biological control,” Bullletin of Entomological Research, vol. 88, no. 4, pp. 461–466, 1998.

[26] J. D. Reeve and W. W. Murdoch, “Aggregation by parasitoids in the successful control of the california red scale: a test of theory,” J Anim Ecol, vol. 54, no. 3, pp. 797–816, 1985.

[27] S. R. Jang and J. L. Yu, “Discrete-time host-parasitoid models with pest control,” J Biol Dyn, vol. 6, no. 2, pp. 718–739, 2012.

[28] M. P. Hassell and G. C. Varley, “New inductive population model for insect and its bearing on biological control,” Nature, vol. 223, no. 1, pp. 1133–1137, 1969.

[29] A. D. Taylor, “Heterogeneity in host-parasitoid interactions: ‘aggregation of risk’ and the ‘*cv*2 > 1 rule.’,” Trends in Ecology and Evolution, vol. 8, pp. 400–405, 1993.

[30] M. P. Hassell, R. M. May, S. W. Pacala, and P. L. Chesson., “The persistence of host-parasitoid associations in patchy environments. I. a general criterion.” American Naturalist, vol. 138, pp. 568–583, 1991.

[31] S. W. Pacala and M. P. Hassell., “The persistence of host-parasitoid associations in patchy environments. II. evaluation of field data.” American Naturalist, vol. 138, pp. 584–605, 1991.

[32] J. D. Reeve, J. T. Cronin, and D. R. Strong., “Parasitoid aggregation and the stabilization of a salt marsh host-parasitoid system,” Ecology, vol. 75, pp. 288–295, 1994.

[33] P. Rohani, H. C. J. Godfray, and M. P. Hassell, “Aggregation and the dynamics of host-parasitoid systems: A discrete-generation model with within-generation redistribution,” The American Naturalist, vol. 144, no. 3, pp. 491–509, 1994.

[34] R. M. May, “Host-parasitoid systems in patchy environments: a phenomenological model,” Journal of Animal Ecology, vol. 47, pp. 833–844, 1978.

[35] P. Rohani and O. Miramontes, “Host-parasitoid metapopulations: the consequences of parasitoid aggregation on spatial dynamics and searching efficiency.” Proc. R. Soc. Lond. B Biol. Sci., vol. 260, pp. 335–342, 1995.

[36] J. T. Cronin and J. D. Reeve, “Host-parasitoid spatial ecology: A plea for a landscape-level synthesis,” Proceedings: Biological Sciences, vol. 272, no. 1578, pp. 2225–2235, November 2005.

[37] F. R. Adler, “Migration alone can produce persistence of host-parasitoid models.” The American Naturalist, vol. 141, no. 4, pp. 642–650, 1993.

[38] H. N. Comins, M. P. Hassell, and R. M. May, “The spatial dynamics of host-parasitoid systems,” Journal of Animal Ecology, vol. 61, no. 3, pp. 735–748, 1992.

[39] B. K. Emerick and A. Singh, “Global redistribution and local migration in semi-discrete host-parasitoid population dynamic models.” bioRxiv, 2019.

